# miR-128 inhibits telomerase activity by targeting TERT mRNA

**DOI:** 10.1101/195198

**Authors:** Herlinda Guzman, Katie Sanders, Adam Idica, Aurore Bochnakian, Douglas Jury, Iben Daugaard, Dimitrios G Zisoulis, Irene Munk Pedersen

## Abstract

Telomerase is a unique cellular reverse transcriptase essential for maintaining telomere stability and required for the unlimited proliferation of cancer cells. The limiting determinant of telomerase activity is the catalytic component TERT, and TERT expression is closely correlated with telomerase activity and cancer initiation and disease progression. For this reason the regulation of TERT levels in the cell is of great importance. microRNAs (miRs) function as an additional regulatory level in cells, crucial for defining expression boundaries, proper cell fate decisions, cell cycle control, genome integrity, cell death and metastasis. We performed an anti-miR library screen to identity novel miRs, which participate in the control of telomerase. We identified the tumor suppressor miR (miR-128) as a novel endogenous telomerase inhibitor and determined that miR-128 significantly reduces the mRNA and protein levels of Tert in a panel of cancer cell lines. We further evaluated the mechanism by which miR-128 regulates TERT and demonstrated that miR-128 interacts directly with the coding sequence of TERT mRNA in both HeLa cells and teratoma cells. Interestingly, the functional miR-128 binding site in TERT mRNA, is conserved between TERT and the other cellular reverse transcriptase encoded by Long Interspaced Elements-1 (LINE-1 or L1), which can also contribute to the oncogenic phenotype of cancer. This finding supports the novel idea that miRs may function in parallel pathways to inhibit tumorigenesis, by regulating a group of enzymes (RT) by targeting conserved binding sites in the coding region of both enzymes.

**NOVELTY AND IMPACT:** Telomerase is an RNA-dependent DNA polymerase that synthesizes telomeric DNA sequences and almost universally provides the molecular basis for unlimited proliferative potential. Expression of human telomerase alone is sufficient for the immortalization of diverse cell types. We have identified the tumor suppressor microRNA (miR-128) as a novel regulator of telomerase, which directly targets the coding sequence (CDS) of TERT mRNA and significantly represses Tert protein expression in a panel of cancer cell lines.

## INTRODUCTION

Limitless replicative potential is considered a hallmark of cancer, which is achieved by an inappropriate reactivation of the essential enzyme telomerase [1, 2]. Telomerase maintains telomere integrity by adding the six-nucleotide repeat sequence, 5’-TTAGGG, to the ends of the chromosomes using its internal template RNA component (TERC) and reverse transcriptase protein component (TERT), thus counteracting the telomere shortening that naturally occurs during DNA replication [3-6]. Telomerase is not expressed in most adult somatic cells and the telomeres are therefore progressively shortened with each round of replication, which ultimately leads to replicative senescence and thus a finite replicative potential. In contrast, stem cells and cancer cells have high telomerase activity. In fact, TERT has been shown to be overexpressed in almost 90% of all human malignancies, but although there is evidence that mutations in the TERT promoter lead to enhanced TERT expression, the mechanisms by which telomerase is reactivated is still poorly understood [7-11].

microRNAs (miRs) are a class of small (~22 nucleotide) non-coding RNAs, which are loaded onto Argonaute (Ago) proteins forming the miR-induced silencing complex (miRISC) and function as post-transcriptional regulators of gene expression by inducing mRNA instability or translational repression [12]. More than 60% of all protein-coding genes are believed to be subjects of miR regulation and alterations in miR expression can therefore have dire consequences and contribute to the development of a wide variety of human diseases, including cancer [12-17]. Depending on their role in carcinogenesis, miRs can generally be divided into oncogenic miRs (oncomiRs) or tumor suppressor miRs that promote or inhibit tumor development and progression, respectively [17-19]. miRs are suspected to be implicated in telomerase reactivation and a subset has been shown to affect telomerase activity directly, *e.g.* by inducing TERT expression. For instance, miR-138 has been reported to function as a direct regulator of TERT expression in thyroid carcinoma and at least 5 additional tumor suppressor miRs (let-7g, miR-133a, miR-342, miR-491 and miR-541) have been shown to be capable of regulating TERT expression through direct interaction with TERT mRNA [20, 21].

Aberrant expression of miR-128 is a frequent observation in human malignancies, but depending on the tumor type, it has been shown to be capable of acting both as an oncomiR and a tumor suppressor miRNA [22, 23]. Most studies have, however, found it to act as a tumor suppressor and downregulation has been documented in a long list of human malignancies, including glioma, prostate, head and neck, lung and colorectal cancer, where it has been shown to function as an inhibitor of cancer cell growth and metastasis [22, 24-28]. We recently demonstrated that miR-128 regulates another cellular reverse transcriptase, namely the Long-Interspaced Element-1 (LINE-1 or L1) by directly interacting with ORF2 L1 RNA, which encodes L1 RT [29]. De-repression of L1 elements have been demonstrated to function as driver mutations during tumor initiation, as well as during tumor progression [30-36].

In this study, we identified miR-128 as a regulator of telomerase activity in an anti-miR library screen, demonstrating that endogenously expressed miR-128 inhibits telomerase activity in HeLa cells. Furthermore, we found that overexpression of miR-128 decreased TERT mRNA and protein levels and miR-128 depletion enhanced the levels of TERT mRNA and protein, relative to controls, in a panel of cell lines. Finally, we demonstrate that miR-128 regulates telomerase activity by directly targeting two sites in the coding region of TERT mRNA. These findings show that tumor suppressor miR-128 also effect the oncogenic phenotype of cancer cells by regulating telomerase.

## RESULTS

### Identification of miR-128 as a regulator of telomerase

We have recently established that miRs (miR-128) can repress the activity of key enzymes in our cells, such as reverse transcriptase (RT) encoded by transposable elements (long-interspaced element-1, LINE-1 or L1) [29]. With this in mind we turned our attention to the most famous RT in human cells – telomerase – an enzyme which plays a crucial role in cancer, stem cells and aging [7-9].

We developed a lenti-viral anti-miR screen for miRs that play a regulatory role of telomerase in HeLa cells. We transduced HeLa cells with an anti-miR library encoding conserved and well-characterized miRs that neutralize the corresponding endogenously expressed miR in the transduced HeLa cells. The lenti-virus also encodes copGFP (green) and puromycin for positive selection of transduced cells. This approach favors a physiologically relevant response by avoiding potential artifacts resulting from ectopic overexpression in cells, which does not normally express a specific miR. Following transduction of the anti-miR library or control miRs, we performed single cell dilutions into 96-well plates and performed a qPCR based functional assay of telomerase activity, using the telomeric repeat amplification protocol (q-TRAP) [9]. The q-TRAP assay involves extension of an oligonucleotide through telomerase-mediated enzymatic addition of telomeric DNA repeats and subsequent PCR amplification of the extension products and serves as a great high throughput functional assay for telomerase activity in cells (Figure 1A).

**Figure 1:**
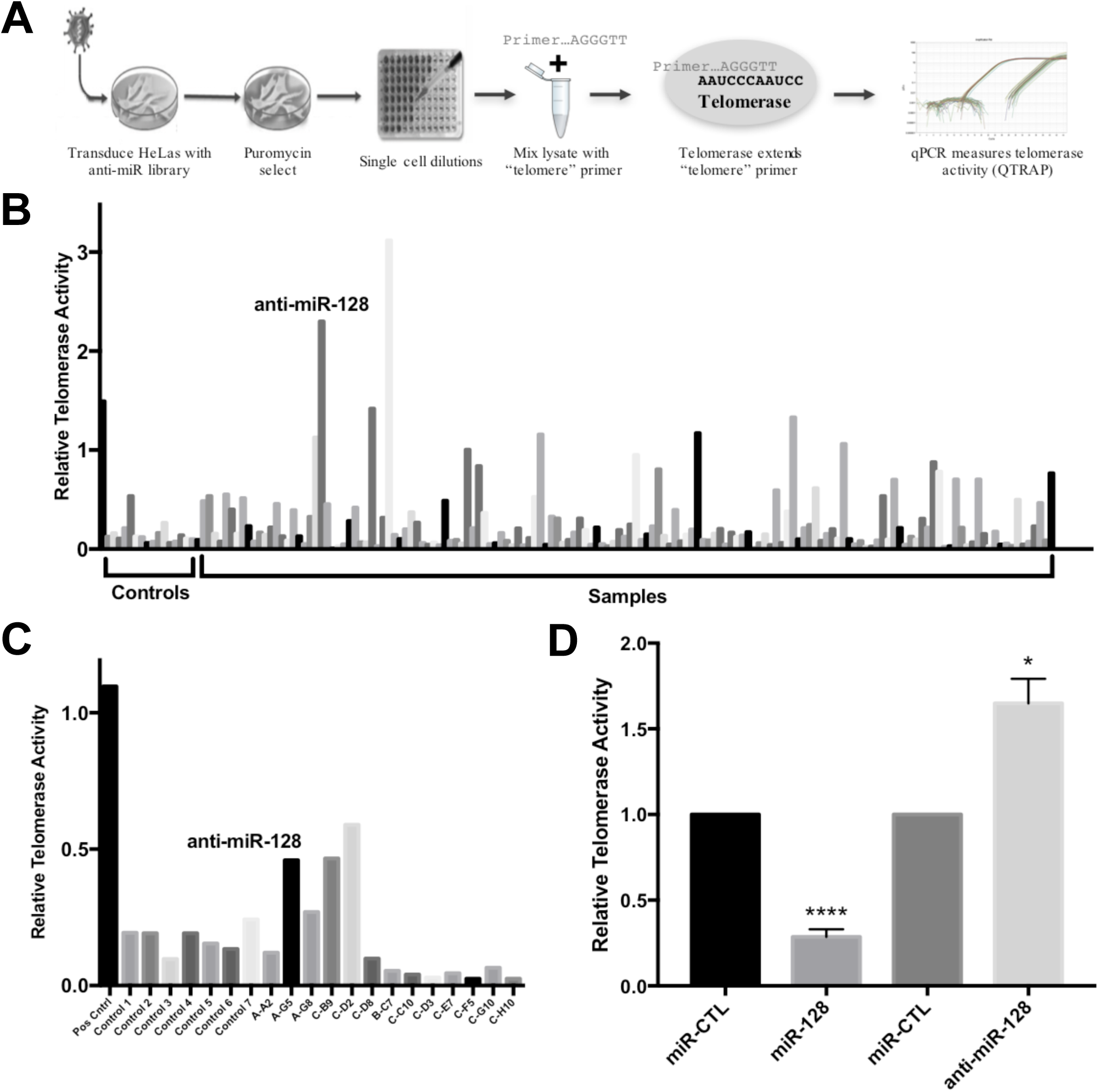
Identification of miR-128 as an inhibitor of telomerase activity. **(A)** Schematic of the anti-miR library screen to identify miRs that regulate telomerase activity. HeLa cells were transduced using a lentiviral-based, miR-neutralizing shRNA library, selected for puromycin resistance and clonally expanded. Each well represents the neutralization of a single endogenously expressed miR. Cells were then subjected to quantitative PCR-based TRAP assay (q-TRAP) analysis using the quantitative telomeric repeat amplification protocol. **(B)** Relative telomerase activity of HeLa cells after transduction with lentiviral miR-neutralizing anti-miR library, selection, and clonal expansion as measured by quantitative telomeric repeat amplification protocol (q-TRAP) as shown. **(C)** Secondary measurement of relative telomerase activity in select samples of anti-miR library-expressing HeLa cells. **(D)** Single high-titer miR-control, miR-128 and anti-miR-128 lentiviruses were generated and stable miR-modulated HeLa cells were assayed for relative telomerase activity measured by q-TRAP analysis. Results shown as percent change ± SEM (n=3, independent biological replicates, *p*<*0.05, ****p<0.0001).

We identified anti-miR-128 as an anti-miR, which significantly de-repress telomeric repeat amplification, as determined by q-TRAP analysis, (Figure 1B). We repeated the assays with a subset of miR modulated HeLa samples and verified that anti-miR-128 enhances telomerase activity (Figure 1C). Next, to evaluate the specificity of the anti-miR-128 effect on telomerase activity, we generated high titer miR-128, anti–miR-128 and control miR lentiviruses, transducing HeLa cells and selecting for puromycin resistance. The miR-128 cell line panel was then evaluated for telomerase activity as described using the q-TRAP assay. As expected anti-miR-128 significantly increased HeLa cell telomerase activity, compared to cells expressing endogenous miR-128 (miR control) (Figure 1D). In contrast, miR-128 significantly reduced the level of telomeric repeat amplification, relative to miR controls (Figure 1D). These experiments suggest that miR-128 regulates telomerase activity in HeLa cells.

### miR-128 reduces TERT mRNA and protein levels

We next wished to characterize the mechanism by which miR-128 regulates telomerase activity. First, we performed RT-qPCR analysis of HeLa cells, a teratoma cell line (Tera-1 or Tera) and an iPSC cell line, in order to determine if miR-128 acts by regulating the amount of TERT mRNA. We found that induced miR-128 expression resulted in a significant decrease in the levels of TERT mRNA in all three cell lines, relative to miR controls (Figure 2A), whereas miR-128 neutralization by anti-miR-128 resulted in an enhanced amount of TERT mRNA. We next examined the potency of miR-128 at regulating Tert protein levels in a small panel of cancer cell lines, including lung cancer (A549), colon cancer (SW620) and pancreatic cancer (PANC1), in addition to HeLa cells (figure 2B). We determined that miR-128 substantially decreased Tert protein levels in all four cell lines, compared to miR controls. In contrast, anti-miR-128 enhanced Tert protein levels, relative to cells expressing miR controls (Figure 2B). Taken together this data demonstrate that miR-128 regulates Tert expression both at the mRNA and protein levels in different cell types.

**Figure 2:**
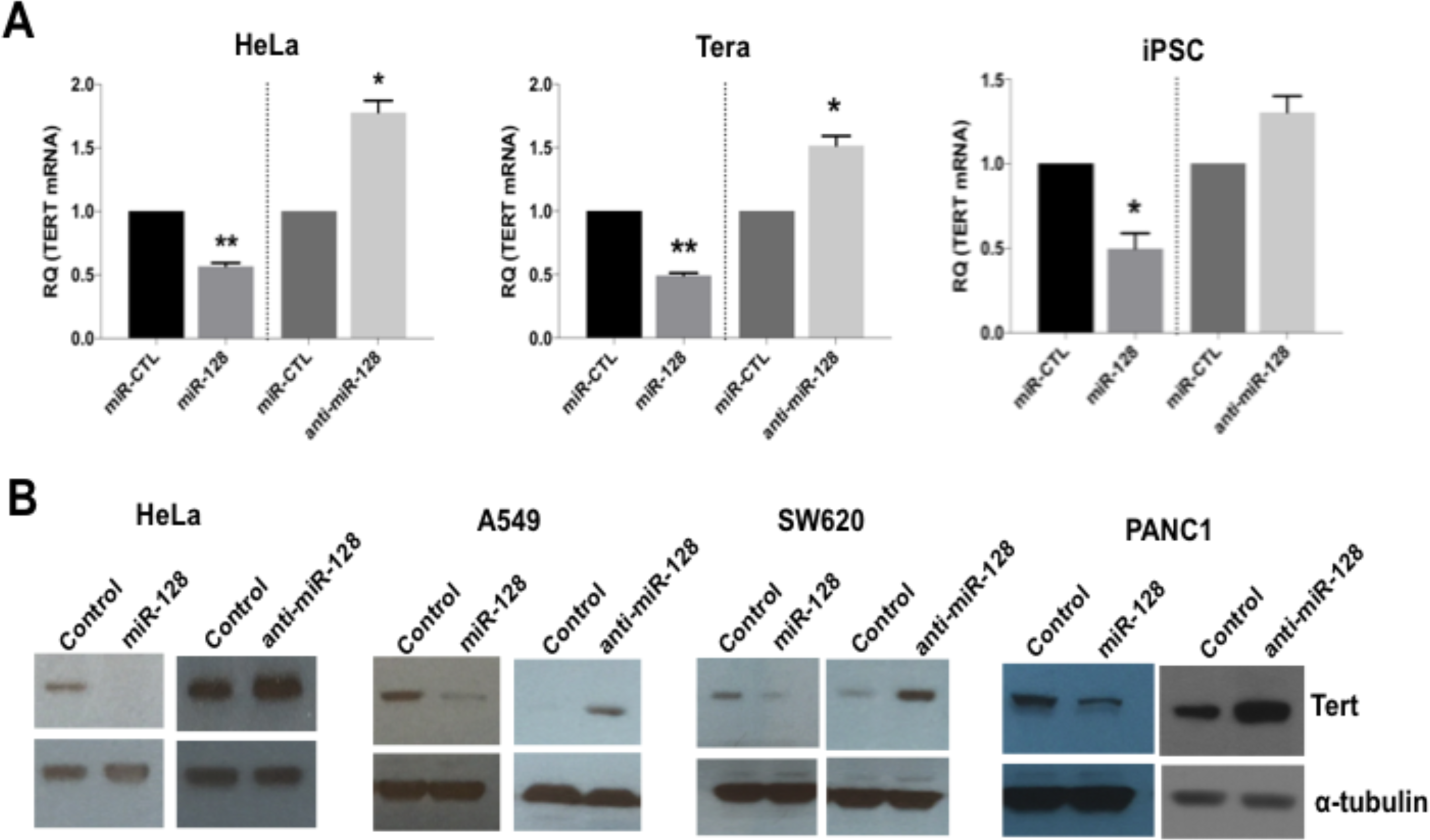
miR-128 reduces the expression of TERT mRNA and protein. **(A)** Stably miR-128, anti-miR-128 or control miR control cell lines were generated of HeLa cell (HeLa), teratoma cell line (Tera) and an induced pluripotent stem cell line (iPSC). Relative expression levels of *TERT* RNA were determined and normalized to beta-2-microglobulin (B2M) expression levels. Results are shown as mean ± SEM normalized to control (n=3 independent biological replicates *p <0.05, **P <0.01). **(B)** Stably miR-128, anti-miR-128 or control miR control HeLa, A549, SW620 ad PANC1 cells lines were generated and western blot analysis of TERT and α-tubulin protein levels were performed (n=3).

### miR-128 interacts with the coding sequence of TERT mRNA

miR-128 could potentially regulate telomerase activity by directly targeting TERT mRNA, or by regulating expression of other proteins that regulate telomerase, or both. In parallel with this work our laboratory have been investigating the mechanisms by which miR-128 regulate L1-induced mutagenesis, including the repression of L1 RT activity. We have demonstrated that miR-128 repress L1 retrotransposition and genomic integration by directly targeting the L1 RNA, in a similar fashion as when miRs function as an anti-viral defense mechanism in human cells, limiting viral replication of RNA virus [29, 37, 38]. We surprisingly determined that miR-128 targets L1 RNA by binding directly through an imperfect seed match to L1 RNA in the coding sequence (CDS) of ORF2, which encodes the L1 endonuclease and reverse transcriptase (RT). With our new finding that miR-128 can also repress telomerase activity in a functional assay (q-TRAP) and by regulating TERT mRNA and protein levels (Figure 2), we hypothesized that miR-128 might be binding to a shared conserved site between L1 RT and TERT mRNA. We therefore aligned the mRNA sequences of the two cellular RTs (L1 and TERT) and determined that the functional miR-128 binding site in L1 is, in fact, present at two locations in the CDS of TERT mRNA (Figure 3A and 3B). To test if the two non-canonical miR-128 seed sites are functional, we generated TERT CDS luciferase constructs either encoding the wildtype (WT) TERT binding Site #1 or binding Site #2. In addition, we generated a 23nt perfect miR-128 match positive control plasmid (as previously described [29, 39]). HeLa cells were co-transfected with one of the three TERT constructs (WT Site #1, WT Site #2 or the positive control) and either miR-128 or miR control mimic oligonucleotides. Luciferase activity was modestly, but significantly reduced in HeLa cells transfected with either one of the two WT TERT constructs and miR-128, relative to miR controls (Figure 3C). As expected, luciferase activity was potently repressed in the positive HeLa cell control (Figure 3C). These experiments supports the conclusion that miR-128 can bind to the predicted binding site likely at both location of the TERT mRNA sequence. Next, we generated a plasmid encoding the TERT binding sites (Site #2) in which we included mutations in the putative miR-128–binding site of TERT mRNA (Figure 3D, top). HeLa cells co-transfected with the plasmid encoding WT TERT and miR-128, showed a significant reduced luciferase activity as previously demonstrated (Figure 3C and 3D). In contrast, HeLa cells co-transfected with the mutant TERT mRNA-binding site (Mut) and either mature miR-128 or control-miR mimics exhibited luciferase activity at similar levels as in the WT TERT and control-miR cells, consistent with miR-128 no longer binding and repressing luciferase reporter-gene expression (Figure 3D). These experiments determine that miR-128 is dependent on the predicted nucleotide sequence to interact with TERT mRNA.

**Figure 3:**
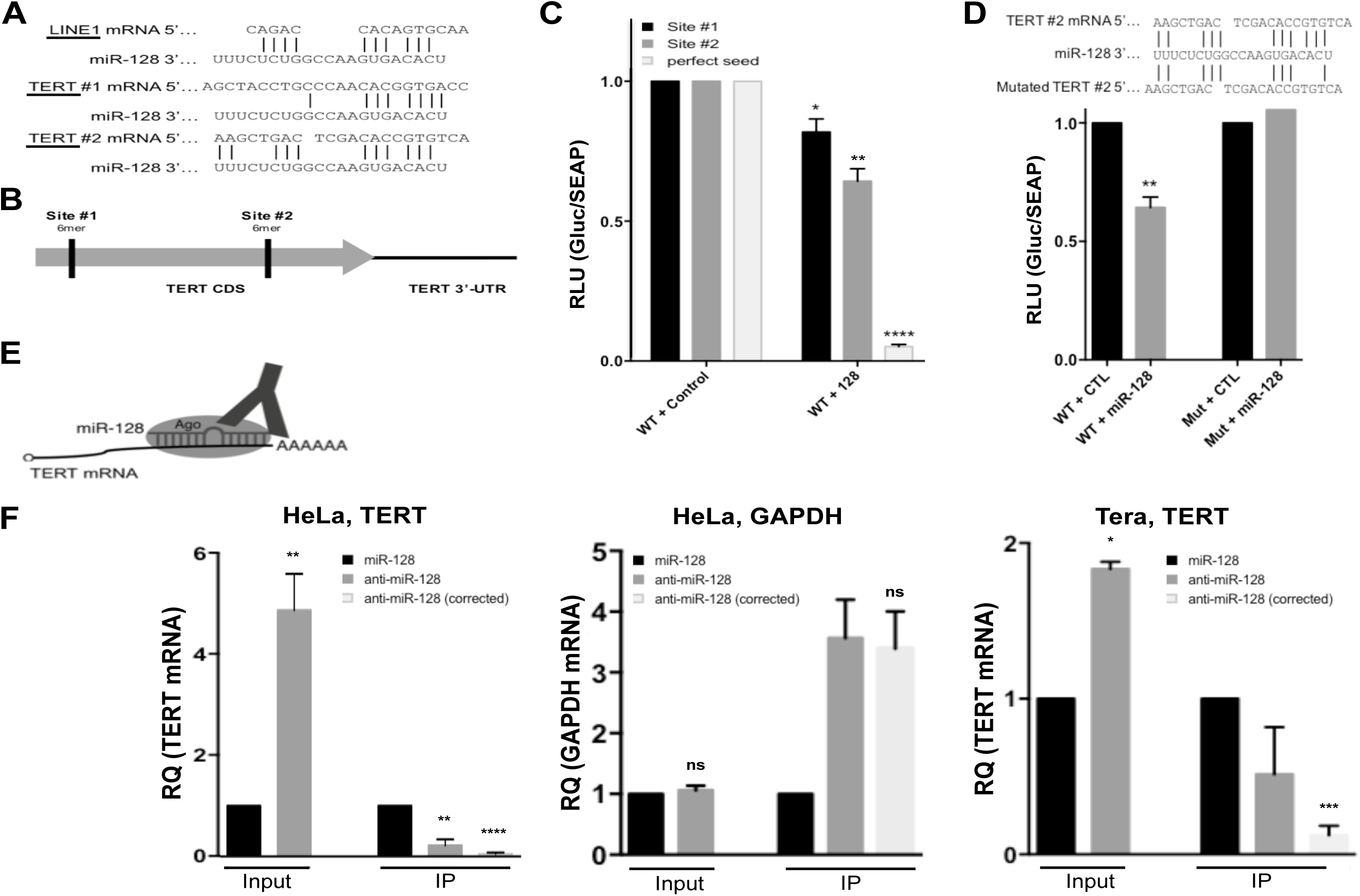
miR-128 directly interacts with the coding sequence of TERT mRNA. **(A)** Schematic representation of the predicted and functional miR-128 binding sites in long-interspaced element-1 (LINE-1, L1) CDS RNA compared to two predicted miR-128 binding sites in the CDS of TERT mRNA. **(B)** Schematic representation of the two putative 6-mer miR-128 binding sites in the coding sequence of TERT mRNA. **(C)** Relative luciferase levels of HeLa cell transfected with constructs expressing either TERT binding sequences at site #1 or site #2 or a perfect miR-128 seed match (perfect seed), along with miR-CTL or miR-128 mimics, measured 48 hrs post transfection. Results shown as percent change ± SEM (n=3 independent experiments, *p<0.05, **P<0.01, ****p<0.0001) **(D)** Relative luciferase levels of HeLa cell transfected with constructs expressing the mutated 6-mer TERT binding sequences, along with miR-CTL or miR-128 mimics, measured 48 hrs post transfection. Results shown as percent change ± SEM (n=3 independent experiments, **p<0.01). **(E)** Schematic of Argonaute immunopurification strategy (Ago-RIP) strategy. HeLa or Tera cell lines were generated in which miR-128 was either neutralized or over-expressed (stably transduced with anti-miR-128 or miR-128-expressing constructs). If TERT is a direct target of the miR-128/Ago complex, then Ago immunopurification in cells with neutralized miR-128 will pull out less TERT mRNA compared to miR-128 expressing cells, which will bind TERT mRNA directly. **(F)** Relative levels of TERT mRNA were determined by q-PCR analysis and (normalized to B2M) in “input samples” of miR modulated HeLa cells. Relative TERT mRNA levels were next determined in IP fractions and normalized to input levels. TERT IP fractions are also shown as “corrected” levels, in which IP TERT levels were corrected for levels of miR-128 in HeLa samples. As a negative control the same samples were analyzed for relative expression of GAPDH levels. Finally, Ago-RIP in Tera cells was performed as described for HeLa cells. Results from 3 independent experiments are shown as mean of IP fraction ± SEM of three independent experiments (n=3, * *p*<0.05, ** *p*<0.01, \*\*\**p*<0.001, **** *p*<0.0001).

Finally, to test whether miR-128 interacts with TERT mRNA in cells, we isolated Ago complexes containing miRs and target mRNAs by immunopurification from HeLa cells either overexpressing miR-128 or anti–miR-128 and assessed relevant complex occupancy by TERT mRNA (Figure 3E). As determined previously (Figure 2A), miR-128 reduces TERT mRNA levels in HeLa cells (Input). In addition, we determined that despite the lower levels of TERT mRNA in HeLa cells (caused by over-expression of miR-128), Ago-bound TERT mRNA was significantly higher in cells overexpressing miR-128 than in cells in which miR-128 was downregulated by anti–miR-128 (IP) (Figure 3F, left panel). When we corrected for the higher levels of TERT mRNA in HeLa cells treated with anti–miR-128, the difference in bound TERT mRNA was even more significant (IP) (anti–miR-128 corrected) (Figure 3F, left panel). A control, constitutively expressed transcript of the GAPDH gene did not show altered levels of total RNA in cells transduced with miR-128 or anti–miR-128 (Input), or relative differences in Ago immunopurification (IP) (Figure 3F, middle panel). Finally, we verified that TERT mRNA could also be immunopurified by miR-128-Ago in Tera cells, showing that miR-128 significantly reduces TERT mRNA levels in Tera cells (Input), and that miR-128 interacts with TERT mRNA in Tera cells, at a significantly manner, when correcting for miR-128 levels (Figure 3F). This body of work supports the conclusion that miR-128 interacts with TERT mRNA and suggests that the putative miR-128 binding site in the coding region of TERT mRNA is indeed the functional binding site resulting in potent regulation of TERT levels and telomerase activity.

## DISCUSSION

This study is the first to identify that miR-128 targets TERT mRNA and reduces TERT mRNA and protein levels resulting in a decrease in telomerase activity in cancer cells. Telomerase is a cellular reverse transcriptase that maintains chromosome health by extending telomeres and protecting chromosome ends. While telomerase is inactive in most adult cells, it is reactivated in cancer cells allowing continuous proliferation. The necessity for telomerase in continued cancer cell growth makes it an attractive therapeutic target.

microRNAs (miRs) have been established as crucial players in cancer initiation and progression by regulating oncogenes or tumor suppressor genes. miR-128 has previously been found to act as a tumor suppressor and it is downregulated in various types of cancer, including glioma [28], lung cancer [40], prostate cancer [25] and bladder cancer [41]. Furthermore, miR-128 is highly enriched in the brain, but not detected in glioma cells. Mechanistically, miR-128 has been found to reduce glioma cell proliferation and promote stem cell self-renewal, by the regulation of the BMI-1 oncogene [28]. miR-128 has also been demonstrated to positively regulate p53 by directly targeting SIRT1, and promote apoptosis in a PUMA-dependent manner [42]. In non-small cell lung cancer cells, miR-128 overexpression was observed to suppress invasion and induce cell cycle arrest and apoptosis. Interestingly, when miR-128 was restored, tumorigenicity was greatly suppressed in a mouse model of lung cancer [43]. Taken together, these studies demonstrates that miR-128 functions as a tumor suppressor in many cancer types and our experimental findings add another mechanism by which miR-128 repress the oncogenic phenotype of cancer cells.

We have previously established that miR-128 surprisingly interacts with the coding sequence (CDS) of the reverse transcriptase component of long interspersed element 1 (LINE1) retrotransposons (ORF2), preventing retrotransposition, genomic integration and mutagenesis [29]. The finding that the functional miR-128 binding sites are located in both the CDS of telomerase mRNA and the ORF2 encoded L1 RT responsible for repressing Telomerase activity and L1 mobilization in cancer cells, is interesting and supports the novel concept that one microRNA (miR-128) can function by parallel regulation of similar cellular enzymes (RTs) by binding to conserved target sequences in the coding region.

## ABBREVIATIONS

Ago: Argonaute
BMI-1: *BMI1* Proto-Oncogene, Polycomb Ring Finger
CDS: coding sequence
GAPDH: Glyceraldehyde 3-phosphate dehydrogenase
GFP: green fluorescent protein
iPSC: induced pluripotent stem cells
L1 or LINE-1: long-interspaced element-1
miR: microRNA
miRISC: miR-induced silencing complex
mRNA: messenger RNA
ORF: open reading frame
PUMA: p53 upregulated modulator of apoptosis
q-TRAP: Real-time quantitative telomeric repeat amplification
RIP: Argonaute-RNA immuno-purification
RT: reverse transcriptase
SIRT1: silent mating type information regulation 2 homolog
shRNA: short hairpin RNA
TERT: Telomerase Reverse Transcriptase
TERC: telomerase RNA component
WT: wild type

## ACKNOWLEDGEMENTS

This work was supported by University of California Cancer Research Coordinating Committee 55205 (IMP), American Cancer Society – Institutional Research Grant 98-279-08 (IMP).

## AUTHOR CONTRIBUTION

KS made the initial identification of miR-128 as a novel telomerase regulator. HG, KS, ES and AI performed the majority of experiments, with the help of DJ and ID, demonstrating that miR-128 targets TERT mRNA and reduces the levels of Tert mRNA and protein. DGZ performed the Ago RNA IPs and IMP directed all experiments, figure design and wrote the manuscript with help from ID.

## MATERIALS AND METHODS

### Cell culture and transductions

All cells were incubated at 37°C and 5% CO_2_ and routinely checked for mycoplasma contamination. HeLa cells (CCL-2, American Tissue Cell Culture (ATCC)) were cultured in EMEM (SH3024401, Hyclone) supplemented with 10% FBS. 293T cells (CRL-3216, ATCC) were cultured in DMEM (25-501N, Genesee) with 10% FBS. Tera-1 cells (HTB-105, ATCC) were cultured in McCoy’s 5A (16600-082, Life Technologies) supplemented with 20% cosmic serum (SH3008702, Fisher Scientific).

### anti-miR library screen

HeLa cells were transduced with miRZip Virus Library (MZIPPLVA, System Biosciences), selected for Puromycin resistance and split to single cell dilutions in 96-well plates. Cells were grown to confluency and telomerase activity was measured using the telomeric repeat amplification protocol (q-TRAP) [9].

### Cell transductions

VSV-G-pseudotyped lentiviral particles were made by transfecting 293T cells with 0.67 µg of pMD2-G (12259, Addgene), 1.3 µg of pCMV-DR8.74 (8455, Addgene) and 2 µg of mZIP-miR-128 or mZIP-anti-miR-128 using Lipofectamine LTX (15338030, ThermoFisher). Viral supernatants were concentrated using PEG-it (LV810A-1, System Biosciences). Cells were transduced with high titer virus using polybrene (sc-134220, Santa Cruz Biotech) and spinoculated at 800 xg at 32°C for 30 minutes. Transduced cells were selected and maintained using 10 µg/ml puromycin.

### Quantitative telomeric repeat amplification protocol (Q-TRAP)

HeLa cells were lysed in NP40 Lysis buffer and q-TRAP analysis was carried out as described previously by Herbert et al. Briefly, lysates were mixed with EGTA (NC9118216), Platinum Taq polymerase (10966034, Life), ACX primer (5’-GCG CGG CTT ACC CTT ACC CTT ACC CTA ACC-3’), TS primer (5’-AAT CCG TCG AGC AGA GTT-3’), and either SYBR Green Master Mix (4367659, Life) or Forget-Me-Not qPCR Master Mix with Rox (31042-1, Biotium). Samples were incubated for 30 min at 30°C for telomerase extension, then 95°C for 10 min to deactivate telomerase and activate Platinum Taq polymerase, then 40 cycles of 95°C for 15 sec and 60°C for 1 min. Amplification was normalized to a standard curve of HeLa lysates diluted 1:5 from 1 µg/well to 8 ng/well.

### RNA extraction and RT-qPCR

RNA was extracted with Trizol (15596018, ThermoFisher Scientific) according to manufacturer’s instructions and cDNA synthesis was performed with the High Capacity Reverse Transcriptase Kit (4368813, Life Technologies). cDNA was amplified relative to GAPDH using the Forget-Me-Not qPCR Master Mix with Rox (31042-1, Biotium) according to the manufacturer’s protocol.

### Western Blot

HeLa cells were lysed in RIPA buffer (89901, ThermoFisher) supplemented with 1x protease inhibitor cocktail (PI78410, ThermoFisher) and then mixed with 4x LDS sample buffer (NP0008, ThermoFisher) and boiled at 95°C for 10 minutes. Samples were run on NuPAGE Novex 4-12% Bis-Tris Protein Gels (NP0335, ThermoFisher Scientific), and transferred to PVDF membranes. Membranes were incubated with rabbit anti-human TERT antibody (1531-1, Epitomics) or rabbit anti-human GAPDH 14C10 antibody (2118, Cell Signaling) then HRP-linked anti-rabbit IgG antibody (7074S, Cell Signaling) and visualized with Pierce ECL Western Blotting Substrate (32106, ThermoFisher) on the Bio-Rad ChemiDoc XRS+ System.

### Luciferase Direct-binding Reporter Assay

Wildtype or mutated TERT sequences were cloned into a dual luciferase reporter plasmid (pEZX-MT05, Genecopoeia). 3 x 10^5^ HeLa cells were forward-transfected with 0.8 µg reporter plasmid and 20 nM control mimic or miR-128 mimic with Attractene transfection reagent (301005, Qiagen) according to the manufacturer’s instructions. Relative Gaussia luciferase and secreted alkaline phosphatase (SEAP) levels were determined with the Secrete-Pair Dual Luminescence Assay Kit (SPDA-D010, Genecopoeia) on a Tecan Infinite F200 microplate reader.

### Argonaute RNA Immunopurifications (AgoRIP)

Immunopurification of Argonaute from HeLa and Tera cell extracts was performed using the 4F9 antibody (4F9, Santa Cruz Biotechnology) as described previously [29]. Briefly, 10 mm plates of 80% confluent cultured cells were washed with buffer A [20 mM Tris-HCl pH 8.0, 280 mM KCl, 10 mM EDTA, 1% NP-40, 0.2% Deoxycholate, 2X Halt protease inhibitor cocktail (Pierce), 200 U/ml RNaseout (ThermoFisher Scientific) and 1 mM DTT]. Protein concentration was adjusted across samples with buffer B [20 mM Tris-HCl pH 8.0, 140 mM KCl, 5 mM EDTA pH 8.0, 0.5% NP-40, 0.1% deoxycholate, 100 U/ml Rnaseout (ThermoFisher Scientific), 1 mM DTT and 1X Halt protease inhibitor cocktail (Pierce)]. Lysates were centrifuged at 16,000xg for 15 min at 4°C and supernatants were incubated with 10-20µg of 4F9 antibody conjugated to epoxy magnetic beads (M-270 Dynabeads, ThermoFisher) for 2 hours at 4°C with gentle rotation. Following magnetic separation, the beads were washed three times five min with 2 ml of buffer C [20 mM Tris-HCl pH 8.0, 140 mM KCl, 5 mM EDTA pH 8.0, 40 U/ml Rnaseout (ThermoFisher Scientific), 1 mM DTT and 1X Halt protease inhibitor cocktail (Pierce)]. Following immunopurification, RNA was extracted using miRNeasy kits (217004, Qiagen), following the manufacturer’s recommendations and qPCR was performed using custom probes/primers and Forget-me-not qPCR master mix (31042-1, Biotium). Results were normalized to their inputs and shown as “corrected” values as a proxy for Ago immunopurification efficiency.

